# Spectroscopic assessment of flavor-related chemical compounds in fresh tea shoots using deep learning

**DOI:** 10.1101/2024.03.05.583504

**Authors:** Lino Garda Denaro, Shu-Yen Lin, Cho-ying Huang

## Abstract

This study employs a deep-learning method, Y-Net, to estimate 10 tea flavor-related chemical compounds (TFCC), including gallic acid, caffeine and eight catechin isomers, using fresh tea shoot reflectance and transmittance. The unique aspect of Y-Net lies in its utilization of dual inputs, reflectance and transmittance, which are seamlessly integrated within the Y-Net architecture. This architecture harnesses the power of a convolutional neural network-based residual network to fuse tea shoot spectra effectively. This strategic combination enhances the capacity of the model to discern intricate patterns in the optical characteristics of fresh tea shoots, providing a comprehensive framework for TFCC estimation. In this study, we destructively sampled tea shoots from tea farms in Alishan (Ali-Mountain) in Central Taiwan within the elevation range of 879–1552 m a.s.l. Tea shoot reflectance and transmittance data (n = 2032) within the optical region (400–2500 nm) were measured using a portable spectroradiometer and pre-processed using an algorithm; corresponding TFCC were qualified using the high-performance liquid chromatography analysis. To enhance the robustness and performance of Y-Net, we employed data augmentation techniques for model training. We compared the performances of Y-Net and seven other commonly utilized statistical, machine-/deep-learning models (partial least squared regression, Gaussian process, cubist, random forests and three feedforward neural networks) using root-mean-square error (RMSE). Furthermore, we assessed the prediction accuracies of Y-Net and Y-Net using spectra within the visible and near-infrared (VNIR) regions (for higher energy throughput and low-cost instruments) and reflectance only (for airborne and spaceborne remote sensing applications). The results showed that overall Y-Net (mean RMSE ± standard deviation [SD] = 2.51 ± 2.20 mg g^−1^) outperformed the other statistical, machine- and deep-learning models (≥ 2.59 ± 2.64 mg g^−1^), demonstrating its superiority in predicting TFCC. In addition, this original Y-Net also yielded slightly lower mean RMSE (± SD) compared with VNIR (2.76 ± 2.41 mg g^−1^) and reflectance-only (2.68 ± 2.74 mg g^−1^) Y-Nets using validation data. This study highlights the feasibility of using spectroscopy and Y-Net to assess minor biochemical components in fresh tea shoots and sheds light on the potential of the proposed approach for effective regional monitoring of tea shoot quality.

## 1. Introduction

Plant metabolism collectively produces many metabolites, crucial in resisting biotic stress and adapting to abiotic pressure. These metabolites also serve as invaluable resources for human health and survival [1–6]. Tea plants are predominantly cultivated in Asia, producing some of the most popular non-alcoholic beverages in the world [7, 8]. The characteristics of tea are primarily composed of its chemical components (tea flavor-related chemical compounds [TFCC] hereafter), including gallic acid (GA), caffeine (CAF) and eight catechin isomers including gallocatechin (GC), epicatechin gallate (EGC), catechin (C), epicatechin (EC), epigallocatechin gallate (EGCG), gallocatechin gallate (GCG), epicatechin gallate (ECG) and catechin gallate (CG) [8]. These chemicals contribute not only to the flavor of a cup of tea but also to active responders to the environments in fresh tea leaves as tea plants grow [9]. The catechins, GA and CAF mainly contribute to the astringency and bitterness of taste, which is the main body of the tea soup, and the catechins oxidation is the essential chemical reaction that determines the tea characteristics during the manufacturing process [10, 11]. The more we understand the TFCC status in tea leaves, the more effectively we can decide on the management of the tea plants and control the quality of tea production.

Methods have been developed to quantify TFCC, such as gas chromatography or high-performance liquid chromatography (HPLC) [12–14]. However, they require pre-treatment, leading to slow data retrieval, and are infeasible for the real-time TFCC assessment. One practical alternative is non-destructive optical (approximately 400–2500 nm) spectroscopy [15, 16]. Previous studies employed optical spectroscopic estimation methods for some TFCC in ground tea leaves [17, 18]. Huang et al. [19] and Wang et al. [20] used leaf reflectance to estimate pigments and four main catechins and CAF in fresh green leaves using partial least squares regression (PLS). However, these linear models may not be suitable for the prediction due to the complex nature of TFCC. Yamashita et al. [6] uses machine learning, including random forests (RF) and cubist, to analyze leaf reflectance and quantify TFCC in fresh tea shoots. However, information may be lost solely relying on leaf reflectance during optimization. Although reflectance provides valuable insights into interactions between reflected photons and tea leaf surface, some TFCC may absorb light at specific wavelengths, and those features may not be able to be delineated by reflectance.

Leaf transmittance, another measurable leaf optical attribute, indicates the proportion of the incident energy passing through a substance without being reflected and absorbed; transmittance variation may also be related to TFCC. By combining both leaf reflectance and transmittance, we could optimize TFCC modeling in fresh tea shoots. Due to the potentially complex relationship between green vegetation optics and TFCC (e.g., [6]), we employed deep learning to derive TFCC from fresh tea shoots in this study. To our knowledge, this has yet to be carried out in previous literature. Deep learning is more effective in handling large and high-dimensional datasets than machine learning, which frequently requires manual feature engineering and domain-specific knowledge to extract relevant features. The key strength of deep learning lies in its ability to automatically learn and extract important features, enabling it to recognize complex patterns and relationships in the data. Hence, the main objective of this study is to use an advanced deep-learning algorithm (Y-Net, a convolutional neural network [CNN] based residual network [ResNet] design approach) to unravel the complex relationship between tea shoot leaf reflectance/transmittance and TFCC.

## 2. Materials and Methods

### 2.1. Tea shoot spectra collection and analysis

We conducted the field campaign in the Greater Alishan (or Ali-Mountain) tea plantation region (23.47 N, 120.69 E) located in Central Taiwan within the elevation range of 800–1600 m a.s.l. (Fig. 1). Fresh tea shoots (mainly the Chin-Hsin-Oolong cultivar [*Camellia sinensis* var. *sinensis*]) were collected from 15 tea farms with the mean (± standard deviation [SD]) elevation of 1225 ± 178 m a.s.l. ranging from 879 to 1552 m a.s.l (Table 1). The data collection period was from April 12 to June 30, 2022, and May 11–24, 2023, encompassing the spring and summer tea harvesting seasons. For each farm, we randomly collected two airtight polyethylene bags of tea shoots, and each bag contained approximately 50–60 samples (an apical bud and three leaves). Tea shoots were placed in a 0 °C cooler right after being destructively sampled and were transferred to the laboratory within the same day (mostly within six hours).

**Figure 1.**
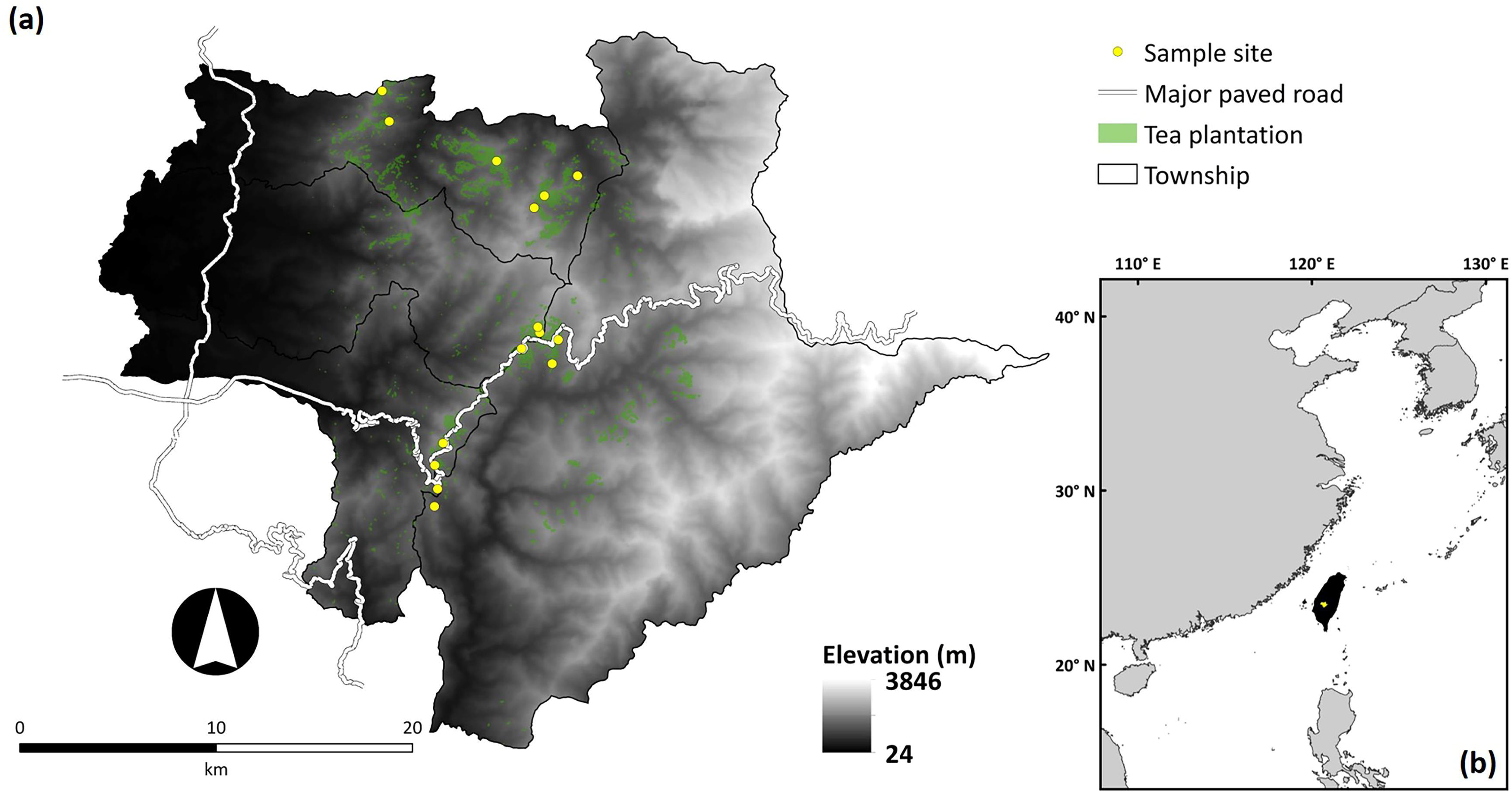
(a) The study area, the Greater Alishan tea plantations (green pixels) within the elevation range of 800–1600 m a.s.l. (b) located in Central Taiwan (the yellow polygon). We collected tea shoot spectral and tea flavor-related chemical compounds data in tea farms (Table 1) illustrated as yellow dots.

**Figure 2.**
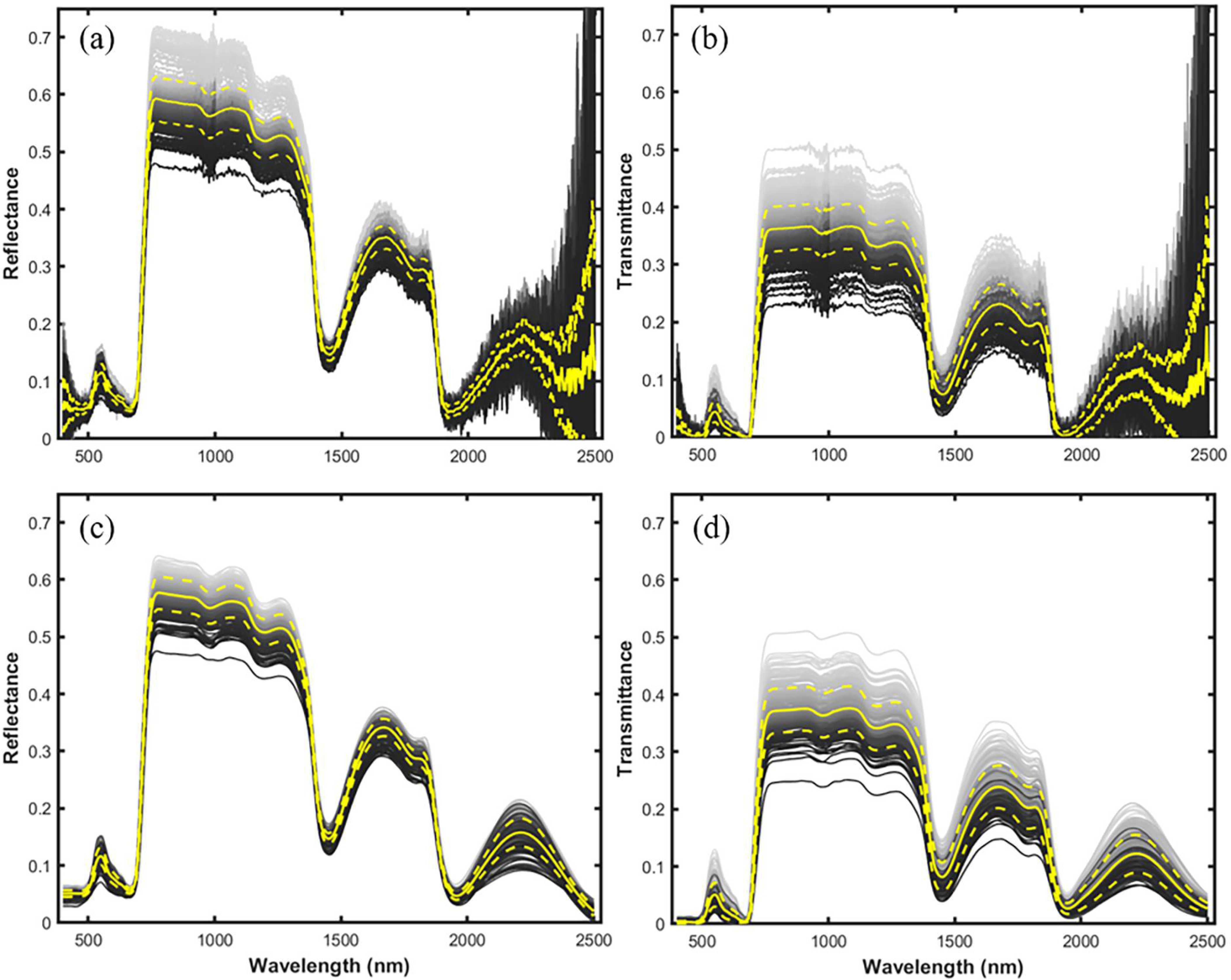
Pre-processing of tea leaf reflectance ((a) before and (c) after the spectral reconstruction) and transmittance ((b) before and (d) after the spectral reconstruction) data.

**Table 1.**
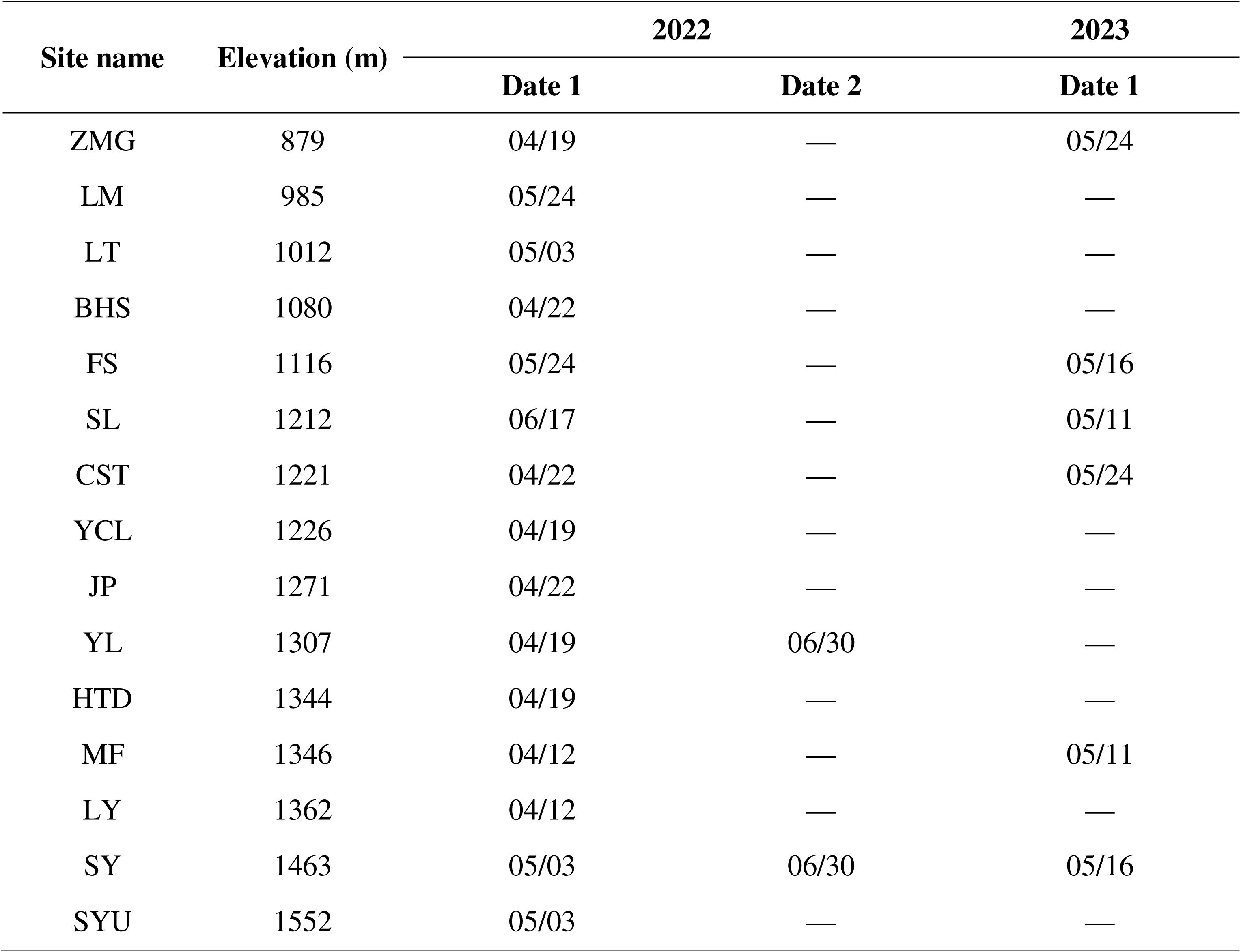
The sample sites and the corresponding elevations (from low to high) and sampling dates. Long dashes (—) indicate no data (dates) available.

We randomly selected 30 samples from each bag to measure tea shoot spectra. Tea leaf reflectance and transmittance were assessed using a portable spectroradiometer (FieldSpec 3, Analytical Spectral Devices, Inc., Boulder, Colorado, USA) with a single integrating sphere (ASD RTS-3ZC). The spectral resolutions were 3 nm (between 350 and 1000 nm) and 10 nm (1000–2500 nm), with sampling intervals set at 1.4 nm and 2 nm, respectively. Due to the sizes of the sample ports of the integrating sphere (diameters ≥ 13 mm), we only measured the spectra of the third leaf (the largest one) for each tea shoot. To ensure the freshness of the tea leaf samples for subsequent biochemical analysis, each spectral reading was an average of 10 measurements. To assess the reflectance (Rs) and transmittance (Ts) of tea leaves, a non-polarized measurement method was employed by referring to the manufacturer-recommended procedure and [21]:

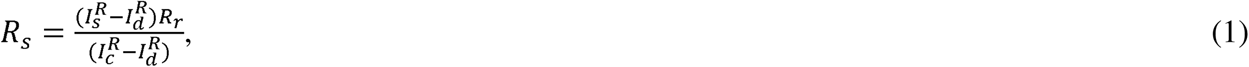

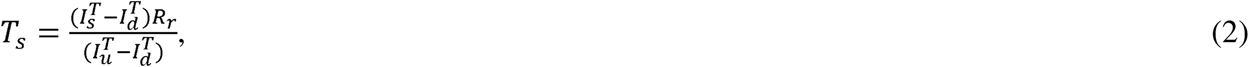

where *I_s_*, *I_c_*, *I_u_*and *I_d_* represent spectroradiometer readings taken from the tea leaf sample, calibrated and un-calibrated white references, and stray light (measured using a light trap), respectively. Superscripts *R* and *T* denote reflectance and transmittance, respectively, and *R_r_* indicates the reflectance factor of the calibrated white reference. The procedure effectively measures the directional reflectance and transmittance factors [22]; the terms “leaf reflectance” and “leaf transmittance” are used for simplicity. One major issue with spectral data collection in a moist region like Taiwan is that high humility may introduce erroneous signals [23, 24]. Hence, we employed a statistical/mathematical spectral reconstruction approach to retrieve noise-free fresh tea leaf spectra involving spectral database matching, multivariable linear regression, linear parameter multiplication and spectral reversion. The procedure is beyond the scope of this study, and details of the spectral pre-processing and reconstruction were described in [25]. There were 1016 pairs (total n = 2032) of reflectance and transmittance data collected in this study.

### 2.2. TFCC analysis

Leaf samples (n = 30) were returned to the polyethylene bags (total 36 bags) after leaf spectral measurement, and the samples were then immediately placed in a −20 °C cold storage unit until TFCC analysis. We note that instead of using the spectrally measured leaf [19], we processed an entire tea shoot (an apical bud and three leaves) since tea makers use the former one to manufacture dried tea products. These samples were freeze-dried, ground into a fine powder, and sifted through a 4 mm sieve. Then, 0.5 grams of this homogeneous tea powder was immersed in 50 ml of boiling deionized water for 20 minutes in a water bath held at 90L. The obtained tea infusions were subsequently filtered via 0.45 μm syringe filters in preparation for HPLC analysis. The HPLC analysis was set up using a Shimadzu system (PU-2089, AS-2057, UV-2075 detector; Shimadzu, Kyoto, Japan) with an equipped C18 column (5 μm × 4.6 mm × 250 mm; Waters, Milford, MA, USA). Solvents A (0.1 % formic acid aqueous solution) and B (the acetonitrile) were introduced following a specific gradient, which involved a linear increment of solvent B from 1% to 10% over 15 minutes, subsequently escalating from 10% to 20% in the next 14 minutes, then further increasing from 20% to 22% within 6 minutes, and lastly maintaining the 22% concentration for an additional 5 minutes. A flow rate of 1.0 ml per minute was maintained throughout the separation process. Absorbance was monitored at 280 nm for real-time tracking of peak intensities. The TFCC components GA, CAF and eight catechin isomers (GC, EGC, C, EC, EGCG, GCG, ECG and CG) were quantified utilizing a calibration curve for each compound, respectively. The calibration curve was formulated using peak areas from HPLC chromatograms at pre-determined concentrations with commercially available chemical compounds (Sigma-Aldrich, St. Louis, MO, USA). All samples were analyzed in triplicate to ensure consistency and accuracy (n = 108).

### 2.3. Deep learning (Y-Net)

#### 2.3.1. Data augmentation

We divided 36 bags of samples into training (70% of the data for developing models) and testing (30% for validation) groups. It is necessary to obtain sizable spectral and TFCC datasets to develop a robust deep-learning model for learning and optimization. However, it is not feasible to produce a large amount of samples of TFCC for deep learning using HPLC due to the nature of the analytical procedure. To mitigate this technical challenge, a simplified approach was adopted using an augmentation process. The rationale for employing the augmentation process stems from the typically distributed correlation between spectral and biochemical characteristics, similar to the principles applied in Gaussian processes [26]. Hence, the augmentation involves using a normal distribution and requires the mean and SD from each bag of TFCC sample to simulate 30 TFCC data to match the sample size of reflectance and transmittance data (illustrated in Fig. 3a). This augmentation technique facilitates expanding the dataset and increases the variation of the available data, thereby enhancing the training process and improving the ability of the model to generalize new data (also see Chen et al. [27] and Wieland et al. [28]).

**Figure 3.**
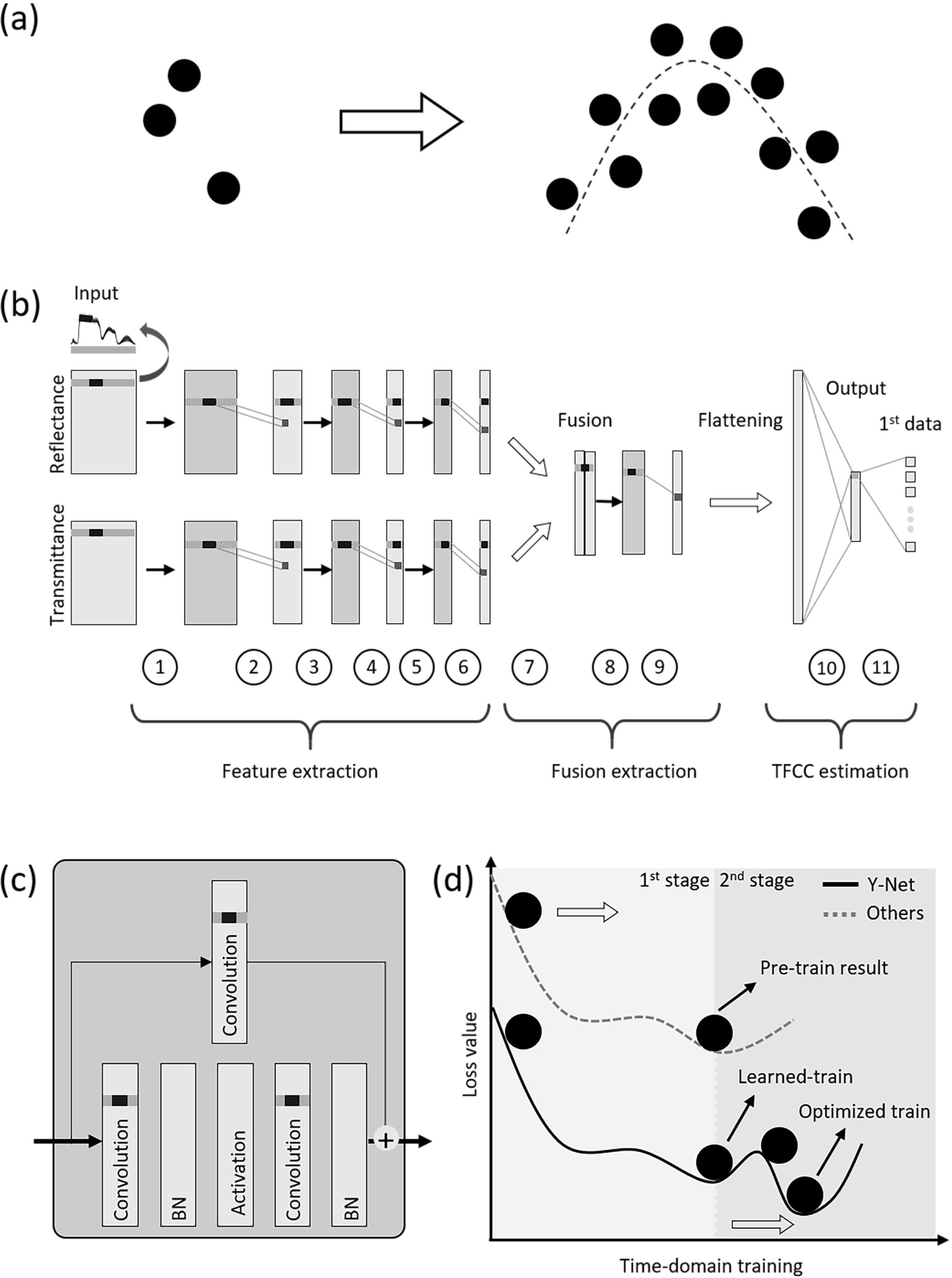
The workflow of using Y-Net to estimate tea flavor-related chemical compounds (TFCC) with tea shoot reflectance and transmittance. (a) Illustration of augmented from observed data. (b) Y-Net-based ResNet (residual network) structure for TFCC prediction. (c) Convolutional block of ResNet, Y-Net. (d) Two-stage training: Top (the first stage) and bottom (the second stage).

#### 2.3.2. Y-Net structure

Y-Net is a specific architecture that utilizes feedforward neural networks (FNN) or CNN-based ResNet approaches. The algorithm can take both reflectance and spectrally corresponding transmittance as input (Fig. 3b). The Y-Net architecture consists of a one-dimensional (1D) convolution block (Fig. 3c), 1D max-pooling, fusion and fully connected layers. The process can be categorized into three steps, including feature extraction, fusion extraction and TFCC prediction:

Feature extraction: This step comprises the encoder, which consists of six processes responsible for extracting meaningful features from the input spectral data. The input dimension was ~×2101×1, indicating one channel and a 1D spectral array with 2101 data points. The symbol ~ represents the number of sample data corresponding to 36 datasets here. The processes numbered 1, 3 and 5 involved convolutional blocks which are part of ResNet processing design, and those with 2, 4 and 6 entailed max-pooling operations (Fig. 3c). The convolutional block operation uses six layers including convolution, batch normalization, activation, and residual layer added to the last layer (Fig. 3d). The kernel size was 1×5, indicating that the convolution operation performed in the spectral windows with a stride of one and using padding to preserve the data dimension. Each convolutional block comprised several kernel filters, and an increased number of kernel filters in each block reduced the spectral dimension. This convolutional block was responsible for feature extraction, mitigating the vanishing gradient problem and preserving information across layers. The max-pooling process occurs between convolutional blocks and was applied to a 1D array using a window size of two. It found the maximum value within each window and applied a stride of two, meaning it moved by two positions at a time. A rectified linear unit function was employed as the activation function in this architecture. It introduced non-linearity by setting negative values to zero and leaving positive values unchanged. The feature extraction step has six convolutional blocks (three reflectance and three transmittance branches). Each corresponding convolutional block (reflectance and transmittance) has a various number of parameters, namely [96, 64, 1296, 64, 96], [2592, 128, 5152, 128, 2592] and [10304, 256, 20544, 256, 10304], in which the total parameters are 10744, and these parameters corresponded to the weights, normalization and biases in the filter masks; where each convolutional block stands for [conv, norm, conv, norm, conv]. These parameters were responsible for capturing and learning the patterns within the data.

Fusion extraction: The fusion extraction consisted of three processes numbered 7, 8 and 9 (Fig. 3c). The fusion process combined the outputs of the right (reflectance) and left (transmittance) processes. It merged the information from these two sources using convolutional block and max-pooling operation, resulting in a fused single processing operation. In the fusion extraction step, one convolutional block fuses the reflectance and transmittance branches. It consists of five parameters [82048, 512, 82048, 512, 82048], which are [conv, norm, conv, norm, conv], respectively. The total number of unknown parameters in the fusion extraction step was 247168. These parameters include normalization, weights and biases associated with the convolutional and max-pooling operations.

TFCC prediction: The process involved in transitioning from the convolutional to the dense processes. This transition included flattening and dropping the data, which converted it from a 2D representation (0-axis for spectral and 1-axis for kernel filters) to 1D dense vectors and passed it through fully connected layers. The reshaped vector had a length of 33290 (Fig. 3c, step 10), which was determined by the size of the output from the last max-pooling in the previous step, which has the dimension ~×26×128. The vector was then connected to a hidden layer with 10 neurons (Fig. 3c, step 11), representing the 10 TFCC components of the first dataset. In the TFCC estimation step, there were 33290 unknown parameters with weights and biases.

A total of 388202 parameters were utilized for optimizing Y-Net, encompassing both the convolutional block and fully connected layers. Studies employed FNNs to determine the optimal neural network structure and parameters, such as filter sizes, number of filters, layers, neurons and activation functions [29, 30]. However, this process often involves trial and error and can be time-consuming. Moreover, due to limited *in-situ* samples and the potential for overfitting, finding the optimal neural structure with the best parameters may not always be feasible. Therefore, this study adopts an alternative strategy by emphasizing the design of the steps within the neural structures rather than exclusively pursuing the optimization of parameters.

#### 2.3.3. Y-Net Transfer Learning

The training of Y-Net was performed in two stages. In the first stage, we applied data augmentation to 1080 samples from 36 batches. In the second stage, the data was transformed by taking the average of three observed data points, resulting in 36 averaged data points corresponding to the 36 batches. Subsequently, we split the data into training (70% of the data) and testing (the rest of the data) sets. The number of the observed points was less than the augmented points to train Y-Net. During Y-Net training, observed data were less than the augmented ones. Hence, there were fewer actual data points available for training Y-Net. Utilizing the augmented data points made it possible to train Y-Net effectively despite the limited number of observed data points. In the initial stage of the two-stage training process (Fig. 3d), the focus was on reaching a local optimum by adjusting the weights, biases and dense parameters from their initial values. The second stage was designed to achieve the global maximum while also reducing the overall training time required to reach the optimal model. In other words, the initial parameters used in the second stage were derived from the pre-trained model obtained during the first stage of training. In the optimization process, we applied the built-in function Adam optimizer. It incorporates adaptive learning rate and moment techniques during training to reach the optimal model. We utilized the mean squared error (MSE) as the loss function (eq. 3):

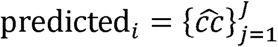

where the 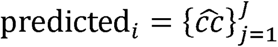 and 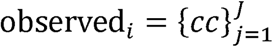; *j* is the index of TFCC *J*, *n* is the number of data, and *cc* and 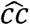 are observed and predicted TFCC, respectively. During the training process, a significant number of epochs were executed and the number of epochs was set to 50. Throughout the training, the parameters of the epoch with the minimal loss value were stored. These optimal parameter values and the network structure were implemented to estimate TCFF.

### 2.4. Performance assessment

To assess the effectiveness and viability of Y-Net, we conducted a comparative analysis by contrasting the performance of the two-stage training with that of the conventional one-stage training. We employed the two-stage training approach to determine whether the additional training stage would enhance model performance by referring to metrics losses. Furthermore, we also selected an advanced multivariate statistical (PLS) approach [31] and other commonly used machine-/deep-learning methods, including cubist [32] with de-trend pre-processing [6], feedforward neural networks (FNNs) [33], Gaussian process [34] and RF [35]. Since these methods have been commonly used in scientific literature, we only provide key references here for expedience. To assess the performances of these methods and Y-Net, we use the root-mean-square error (RMSE) as the evaluation metric, as it measures the average deviation between the predicted and actual observed values. To further evaluate the consistency and stability of the models, we also took the mean and SD into account.

### 2.5. Y-Net spectral analysis

It is important to understand the contribution of different tea shoot spectral regions to estimate TFCC in Y-Net. Therefore, we carried out a spectral feature importance analysis. The feature important analysis took the features of each hidden layer in reflectance and transmittance branches. From the first layer of each branch to the fusion layers, we resampled, averaged and normalized those to the scale of 0–1 to assess their importance to TFCC prediction. The feature importance analysis can be categorized into the local and global aspects to identify the pattern. The global pattern constitutes a significant contribution that can be visually identified, whereas the local pattern represents a minor contribution that requires a tool to unveil subtle information. Therefore, the high-pass filter was also adopted using a fast Fourier transform to enhance the minor contribution by capturing the high frequency in the feature importance [36]. In addition, we carried out a spectral subset analysis. We investigated if visible and near-infrared spectral regions (VNIR, 400–1100 nm) would be sufficient to estimate TFCC using Y-Net since they are responsive to photosynthesis/non-photosynthesis pigments (mainly chlorophyll a and b, carotenoids, xanthophylls and anthocyanins) [37] and cell structure (e.g., the mesophyll cells and the fraction of air space) [38, 39] of fresh green leaves. Therefore, VNIR could be much more sensitive than leaf water content dominant shortwave-infrared bands (SWIR, about 1300–2500 nm) [40]. Moreover, the spectroscopic measurement may be preferable within VNIR due to the relatively higher energy throughput by referring to Planck’s radiation law and more cost-effective for the instruments (silicon/InGaAs for VNIR vs. InSb). Finally, we used tea shoot reflectance of the entire optical region only (without transmittance) for Y-Net since the physical property may be measured by airborne or spaceborne sensors permitting regional mapping of TFCC.

## 3. Results

### 3.1. Input data assessment

To evaluate the performance of Y-Net, it is important to compare the data distribution of the observed data and augmented data (Fig. 4). We found that they have similar characteristics by referring to statistics indicating the central tendency and spread of the data (Table S1). Therefore, the observed and augmented data have similar distributions, which justifies using the latter for further analysis. In addition, to investigate the data distribution of observed and augmented data, a loss value (MSE) comparison for a hundred epochs was conducted in the optimization process (Fig. 5). The optimization process provided two stages of training, with the first stage using augmented data and the second stage using observed data. There was an apparent improvement in the second stage, especially at the initial epoch; the difference in MSE became relatively negligible after 25 epochs. The performance of the two-stage training through this optimization was satisfactory by comparing the observed and predicted training and validation data (Fig. 6 and Table S2). Overall, the data distributions of those groups were comparable except for some outliers of C, ECG and CAF.

**Figure 4.**
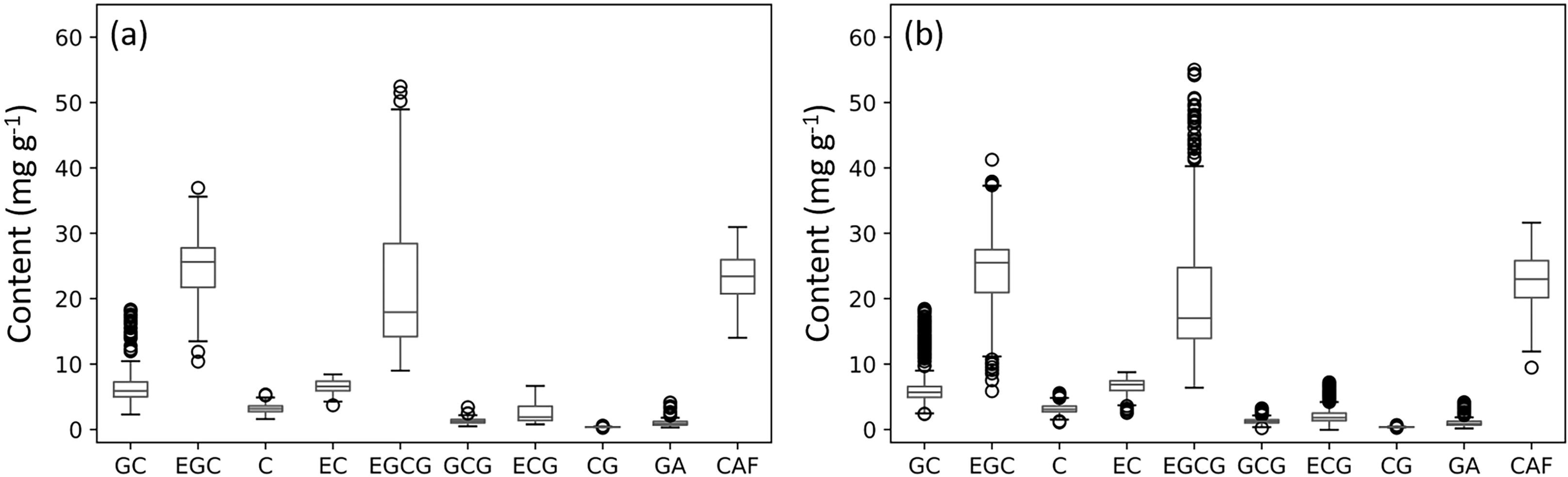
The (a) observed (n = 114) and (b) augmented (n = 1135) TFCC including gallic acids (GA), caffeine (CAF) and eight catechin isomers including gallocatechin (GC), epicatechin gallate (EGC), catechin (C), epicatechin (EC), epigallocatechin gallate (EGCG), gallocatechin gallate (GCG), epicatechin gallate (ECG) and catechin gallate (CG).

**Fig. 5.**
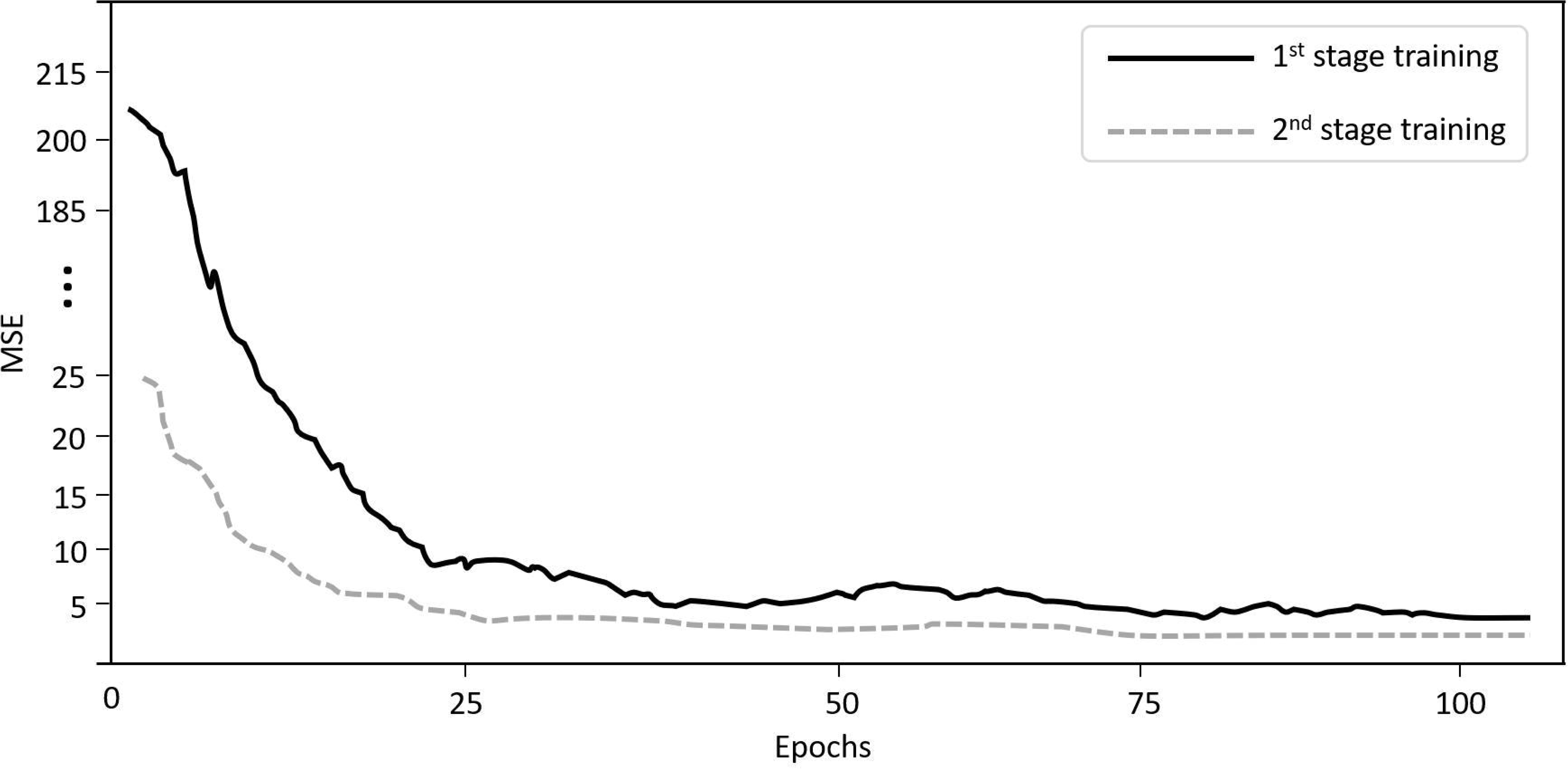
Epoch comparison for 1^st^ and 2^nd^ stage learning; MSE represents the mean squared error.

**Fig. 6.**
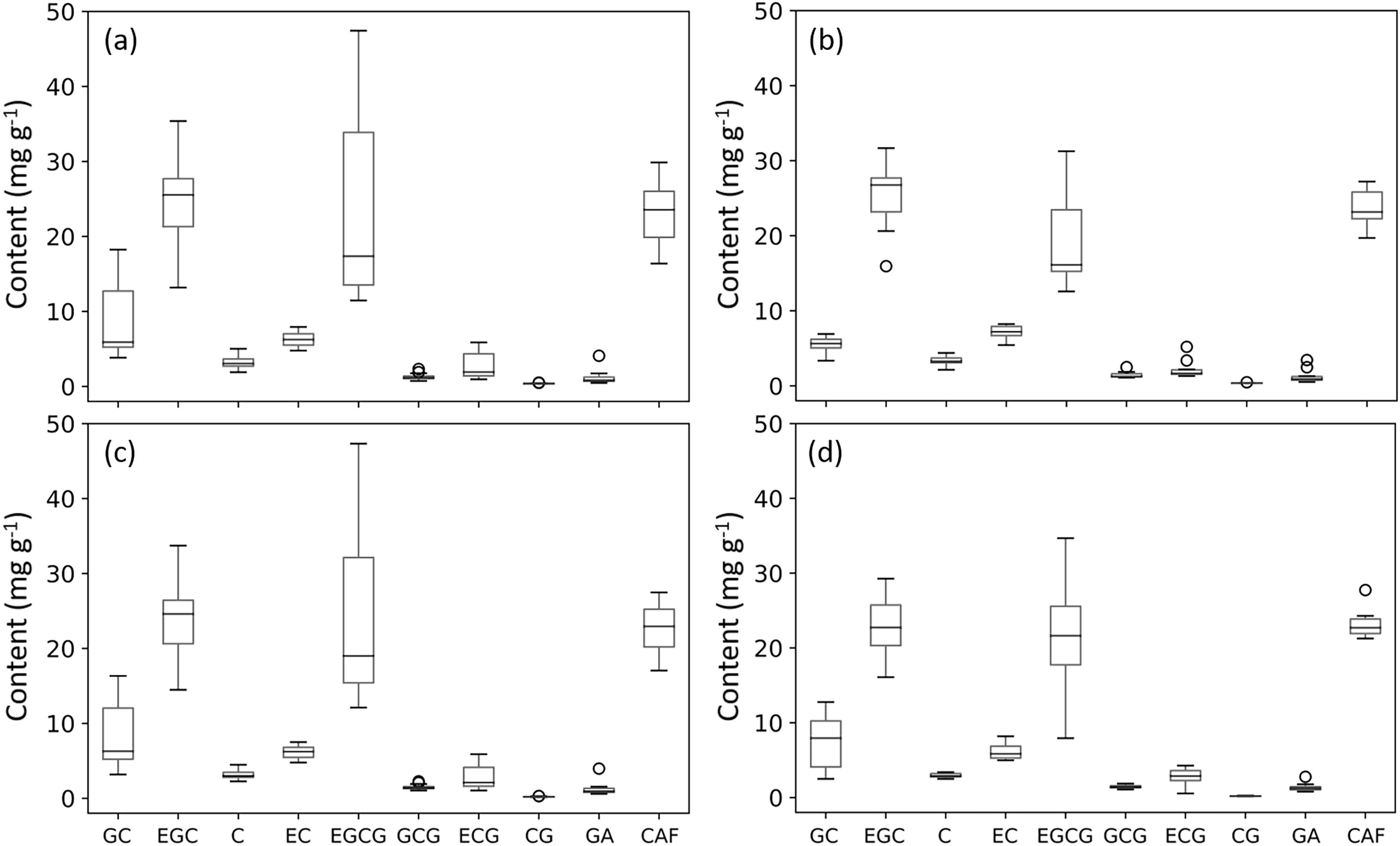
Estimation of TFCC using Y-Net with tea shoot spectra: Observed (a) training and (b) validation; predicted (a) training and (b) validation datasets.

### 3.2. Model performance comparison

Y-Net underwent pre-training and acquired knowledge through a novel two-stage training approach (Fig. 5), utilizing TFCC from observed and augmented data (Fig. 4 and Table S1). Overall, the performance of Y-Net (mean ± SD of RMSE = 2.51 ± 2.20) to model the variation of TFCC was superior to (yielding the lowest mean and SD) selected statistical (PLS) (7.86 ± 7.58) and machine learning (cubist, FNNs, Gaussian, RF) methods (2.59 ± 2.64). The TFCC assessment of Y-Net was relatively consistent by referring to SD demonstrating its effectiveness across various compounds. However, the performance for modeling each TFCC compound differed (Table 2).

**Table 2.** Performance comparison of selected statistics (partial linear least squared [PLS]) and machine-/deep-learning methods (cubist, three feedforward neural networks [FNN-1, FNN-2 and FNN-3], Gaussian process, random forests [RF] and Y-Net based convolutional neural network) to estimate TFCC using root-mean-square error (RMSE). The underscored numbers indicate the lowest values for each TFCC and the mean and standard deviation (SD).

| TFCC | PLS | Cubist | FNN-1 | FNN-2 | FNN-3 | Gaussian | RF | Y-Net |
| --- | --- | --- | --- | --- | --- | --- | --- | --- |
| GC | 15.03 | 8.33 | 3.38 | 3.35 | 3.31 | <u>1.24</u> | 3.74 | 3.23 |
| ECG | 9.02 | 4.33 | 7.93 | 7.92 | 8.57 | 21.27 | <u>5.71</u> | 8.03 |
| C | 4.34 | <u>0.63</u> | 0.74 | 0.73 | 0.83 | 1.55 | 0.57 | 0.86 |
| EC | 2.68 | 1.29 | 1.53 | 1.52 | 1.59 | 2.63 | <u>1.14</u> | 1.49 |
| EGCG | 25.28 | 6.28 | 8.13 | 8.05 | 6.88 | 16.12 | 9.41 | <u>4.09</u> |
| GCG | 1.63 | 0.51 | <u>0.28</u> | <u>0.28</u> | <u>0.28</u> | 3.31 | 0.48 | 0.40 |
| ECG | 4.52 | <u>1.13</u> | 1.52 | 1.5 | 1.24 | 2.86 | 1.68 | 2.24 |
| CG | 0.12 | 0.3 | 0.06 | <u>0.05</u> | <u>0.05</u> | 4.37 | <u>0.05</u> | 0.36 |
| GA | 2.10 | <u>0.8</u> | 0.95 | 0.94 | 0.98 | 3.60 | 0.94 | 1.24 |
| CAF | 13.92 | <u>2.37</u> | 2.79 | 2.83 | 3.31 | 19.31 | 2.77 | 3.19 |
| Mean | 7.86 | 2.59 | 2.73 | 2.72 | 2.70 | 7.63 | 2.65 | <u>2.51</u> |
| SD | 7.58 | 2.64 | 2.82 | 2.81 | 2.75 | 7.52 | 2.80 | <u>2.20</u> |

### 3.3. Y-Net spectral analysis

According to our spectral feature importance analysis, several spectral regions of reflectance and transmittance were pivotal for modeling TFCC; reflectance (55% of the optical spectral bands) played a more critical role than transmittance (45%) by referring to feature important values (Fig. 7). Specifically, after applied the high-pass filter (adopted with fast Fourier transform), spectral bands of reflectance and transmittance located around 400 nm, 700 nm, 1400 nm, 1900 nm and 2500 nm were identified to be crucial for indirectly estimating TFCC. The spectral subset analysis depicts that, overall, the Y-Net using both reflectance and transmittance of the entire optical region (400–2500 nm) (mean ± SD of RMSE = 1.29 ± 1.08 for training and 2.51 ± 2.20 for validation datasets) (Table 3) performed slightly better and relatively more consistent (with the lowest mean and SD) than the VNIR (2.72 ± 2.49; 2.76 ± 2.41) and the reflectance-only (2.07 ± 2.11; 2.68 ± 2.74) Y-Nets. However, the performance of estimating each TFCC compound was different.

**Figure 7.**
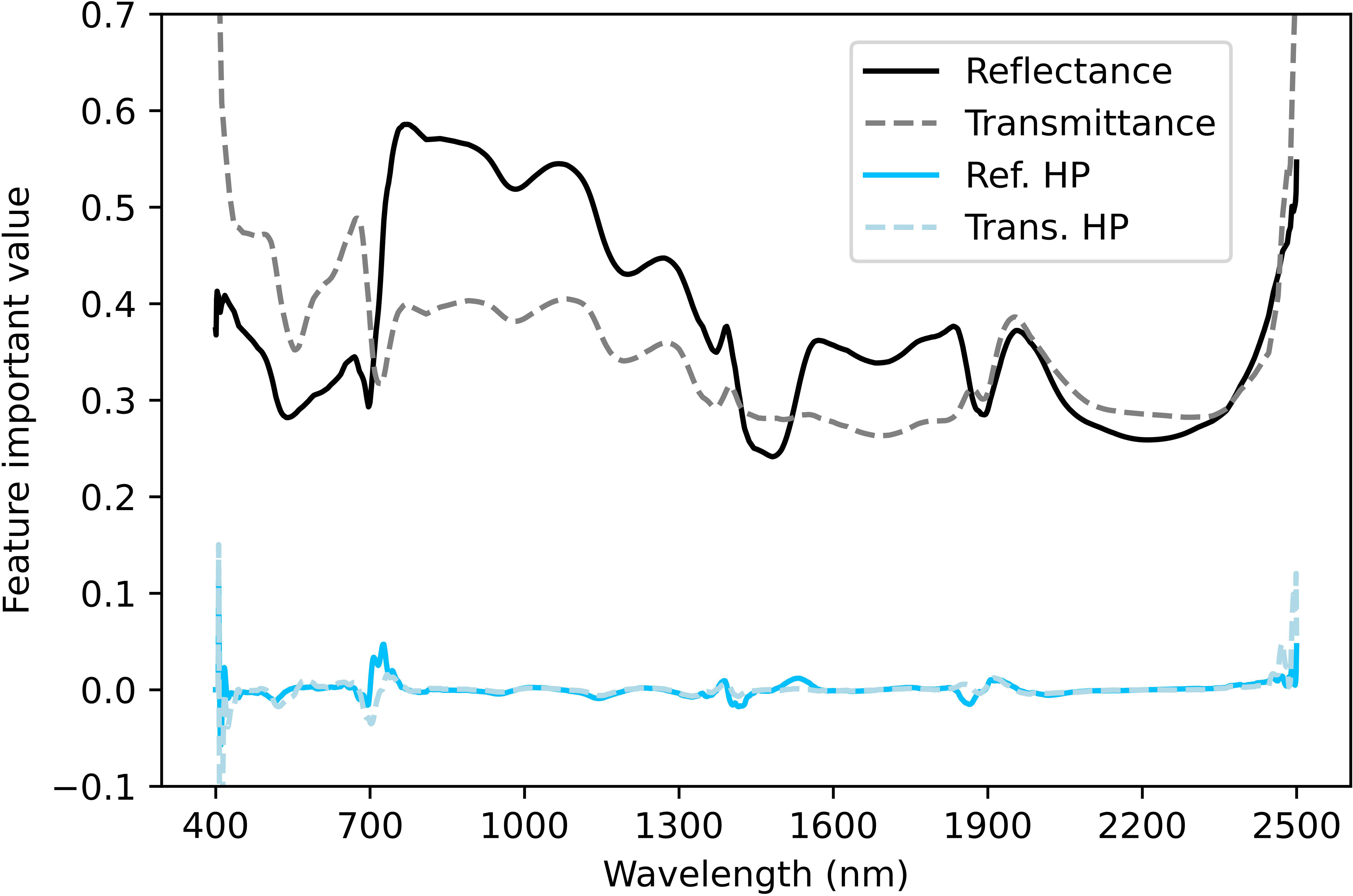
Feature important values of tea shoot optical spectra in Y-Net.

**Table 3.** Y-Net performance comparison (RMSE) using spectra (reflectance and transmittance) from the whole optical (400–2500 nm) and visible-near infrared (400–1100 nm) regions and reflectance of the optical region only to estimate TFCC. The underscored numbers indicate the lowest values for each TFCC, mean and SD, separated by training and validation.

| TFCC | Optical region |  | VNIR |  | Reflectance |  |
| --- | --- | --- | --- | --- | --- | --- |
|  | Training | Validation | Training | Validation | Training | Validation |
| GC | <u>1.57</u> | 3.23 | 2.88 | <u>2.00</u> | 2.75 | 2.98 |
| ECG | <u>2.69</u> | 8.03 | 4.64 | <u>5.63</u> | 4.16 | 7.94 |
| C | <u>0.47</u> | 0.86 | 0.96 | 1.19 | 0.64 | <u>0.70</u> |
| EC | <u>0.66</u> | <u>1.49</u> | 1.43 | 2.02 | 0.92 | 1.70 |
| EGCG | <u>3.57</u> | <u>4.09</u> | 6.64 | 7.90 | 7.12 | 7.70 |
| GCG | <u>0.30</u> | 0.40 | 0.39 | 0.40 | 0.37 | <u>0.34</u> |
| ECG | <u>0.63</u> | 2.24 | 1.61 | 2.42 | 1.12 | <u>1.72</u> |
| CG | 0.38 | 0.36 | <u>0.25</u> | <u>0.29</u> | 0.38 | 0.36 |
| GA | <u>0.54</u> | 1.24 | 0.95 | <u>0.88</u> | 0.74 | 1.21 |
| CAF | <u>2.03</u> | 3.19 | 7.43 | 4.87 | 2.90 | <u>2.79</u> |
| Mean | <u>1.29</u> | <u>2.51</u> | 2.72 | 2.76 | 2.07 | 2.68 |
| SD | <u>1.08</u> | <u>2.20</u> | 2.49 | 2.41 | 2.11 | 2.74 |

## 4. Discussion

### 4.1. Y-Net performance

The Y-Net architecture features a dual-branch input, integrating a ResNet-based CNN [41, 42]. The dual input of Y-Net involves data augmentation or enrichment and pre-training through a novel two-stage training approach (Fig. 3a and 3d), including utilizing TFCC obtained from both augmented and observed data measurements. This unique process markedly improves the performance in predicting TFCC, as evidenced by the initial epoch of the second-stage training involving pre-training using augmented data. Compared to conventional machine-learning methods, the enhancement in performance is demonstrated (see Table 2). The reduction in loss during this phase can be attributed to the utilization of pre-trained parameters from the first-stage training in the second stage. This process serves a dual purpose: Preventing overfitting arising from limited observational data and augmenting the capacity for model generalization. Furthermore, this approach reinforces the adaptability and robustness of the model. It facilitates effective generalization of model performance across diverse datasets, utilizing data augmentation and chemical compounds (TFCC) through feature extraction, fusion and estimation (Fig. 3b). Therefore, Y-Net becomes adept at capturing intricate data patterns with a specific focus on optimizing both local and global aspects during training. The local aspect of Y-Net involves capturing details or specific patterns of the input data. In contrast, the global aspect recognizes the overarching trend or general pattern spanning the entire dataset. Additionally, integrating ResNet into the architecture addresses challenges such as vanishing gradients and ensuring information preservation across layers (Fig. 3c and 3d).

### 4.2. Spectral importance to TFCC retrieval using Y-Net

To assess the performance of Y-Net, we aim to compare the spectral effectiveness of the optical region (400–2500 nm) using feature important values by calculating the average feature extraction values in each branch, namely reflectance and transmittance. The average importance was derived from each convolutional block in the deep CNN (total of 37 layers). Each layer can yield results for feature importance, referred to as spectral contribution to predict TFCC. In the hidden layers, higher spectral feature values contribute significantly to the prediction of TFCC, whereas lower values indicate a comparatively less influential role in the predictive process. The analysis of spectral contributions across layers aids in understanding the significance of various features and their impact on accurately estimating TFCC concentrations by Y-Net throughout the optical range, particularly at 400 nm, 700 nm, 1400 nm, 1900 nm and 2500 nm (Fig. 7). To qualitatively identify patterns, encompassing both local and global aspects, and to isolate local patterns within the branch layers of reflectance and transmittance, a high-pass filter was employed in this study [43, 44]. The high-pass filter adopts a fast Fourier transform that can enhance the minor contribution or local aspect by capturing the high frequency in the feature importance [35]. It is acknowledged that global aspects include visually apparent patterns such as trends and curves of the feature importance. Those global patterns are a significant contribution to TFCC prediction. On the other hand, interpreting local aspects may be challenging due to their minor contribution despite utilizing a substantial influence on TFCC prediction within specific wavelength regions. Using the high-pass filter, we can extract local patterns instead of considering them as interference, especially the feature importance in the deep layer networks. The local pattern was exhibited and appeared in the wavelength at around 400, 500, 700, 1150, 1400, 1900 and 2500 mm (Fig. 7). These significant patterns are evident at the two ends of the optical region (400 nm, 2500 nm) and at 1900 nm, while lower significances were observed around 1150 nm and 1400 nm.

We also compared the spectral effectiveness of the optical region (reflectance and transmittance) with the models ingesting the VNIR range of 400–1100 nm of reflectance and transmittance (due to high signal-to-noise ratio and cost-effective instrument), and reflectance only (for airborne and spaceborne remote sensing applications) of the entire optical region (Table 3). Since the spectral patterns of green vegetation reflectance and transmittance are similar [38] (Fig. 2), they yielded almost identical feature important values (Fig. 7). Thus, the reflectance method was adopted as the comparison in this study. The comparison was visually addressed using the feature importance and their high-pass filter, specifically for reflectance and transmittance, in each branch, while the transmittance was not used in the reflectance method. This implies that by employing VNIR in the experiment, we neglect the significant important features and the less significant features in the spectral range of 1101 nm to 2500 nm (Table 3, VNIR), namely that evident in the spectral edges (400 nm, 2500 nm) and at 1900 nm. Therefore, the proposed Y-Net using entire optical region reflectance/transmittance outperformed the other two Y-Nets models. This is because, although they share almost identical important features and local patterns, they complement each other, with specific patterns present in spectral reflectance that may not be found in spectral transmittance. We also note that the performance of the Y-Net was merely slightly better than the VNIR and reflectance-only Y-Nets. The spectral analysis demonstrates that we could re-produce similar outcomes with Y-Net using low-cost instruments. In addition, this study also underscores the potential of using airborne or spaceborne hyperspectral remote sensing for regional monitoring of TFCC.

### 4.3. Uncertainties and limitations

While Y-Net demonstrates satisfactory performance (Table 2), uncertainties exist, primarily attributed to the complexity of TFCC. The intricate interactions and dependencies among various TFCC pose challenges in achieving absolute precision. The performance of Y-Net is contingent on the quality and diversity of the training data, such as using additional augmented data (Fig. 3a). Limited representation of specific chemical profiles in the training set may lead to challenges in accurately predicting less-represented compounds. For instance, the range of chemical compound values in augmented data may not be sufficient for effective Y-Net training. Expanding the range of compound values in Y-Net involves optimizing sensitivity and precision. Additionally, the generalization of Y-Net to diverse tea cultivars and physical environments could pose a limitation. The inherent variability in TFCC across different varieties and growth environments could ramify the predictive accuracy. Continuous refinement and expansion of the training dataset with actual data (without data augmentation) are crucial measures that could address some of these limitations over time. This ongoing effort aims to enhance the adaptability and robustness of Y-Net to a broader spectrum of TFCC profiles, ultimately improving its applicability across diverse environmental conditions and TFCC.

## 5. Conclusions

In this study, we proposed Y-Net, a deep-learning approach, to estimate TFCC from fresh tea shoots indirectly using their optical reflectance and transmittance. We found that the performance of Y-Net was superior to some commonly employed advanced statistical (PLS) and machine-/deep-learning (cubist, FNNs, Gaussian and RF) methods by referring to RMSE. The dual-input approach of Y-Net, considering both reflectance and transmittance simultaneously, outperformed those selected single-input models, providing a more comprehensive representation of TFCC and improving predictive accuracy. Our spectral analyses showed that some specific bands located at 400 nm, 500 nm, 700 nm, 1150 nm, 1400 nm, 1900 nm and 2500 nm were important for retrieving TFCC. In addition, the reduction of the spectral range (VNIR Y-Net) or exclusion of transmittance (reflectance only Y-Net) would also retard the overall performance. These results emphasize the importance of advanced optimization techniques and leveraging multiple sources of spectral data for precise chemical compound prediction in fresh tea shoots.

## Acknowledgments

We appreciate the field and lab assistance provided by Chi-Ching Huang, Yi-Ching Hung and Zih-Yu Shen. This work was supported by the National Science and Technology Council (NSTC 112-2321-B-002-016), National Taiwan University (NTU-AS-112L104303), and the Research Center for Future Earth, the Featured Areas Research Center Program, the Higher Education Sprout Project, Ministry of Education (Taiwan).

