## Supplemental data for "Spectroscopic assessment of flavor-related chemical compounds in fresh tea shoots using deep learning"

**Table S1.** Statistics (mean and standard deviation [SD]) of observed and augmented TFCC data including gallic acids (GA), caffeine (CAF) and eight catechin isomers including gallocatechin (GC), epicatechin gallate (EGC), catechin (C), epicatechin (EC), epigallocatechin gallate (EGCG), gallocatechin gallate (GCG), epicatechin gallate (ECG) and catechin gallate (CG).

| **TFCC** | **Observed data** | | | | **Augmented data** | | | |
| --- | --- | --- | --- | --- | --- | --- | --- | --- |
|  | **Mean** | **SD** | **Skewness** | **Kurtosis** | **Mean** | **SD** | **Skewness** | **Kurtosis** |
| GC | 6.59 | 3.29 | 2.31 | 4.94 | 6.60 | 3.31 | 2.27 | 4.51 |
| ECG | 24.48 | 5.26 | -0.25 | -0.09 | 24.47 | 5.25 | -0.24 | 0.16 |
| C | 3.19 | 0.70 | 0.45 | 0.54 | 3.19 | 0.70 | 0.46 | 0.51 |
| EC | 6.63 | 1.09 | -0.50 | -0.58 | 6.62 | 1.11 | -0.55 | -0.36 |
| EGCG | 20.08 | 8.71 | 1.34 | 1.45 | 20.04 | 8.54 | 1.28 | 1.32 |
| GCG | 1.31 | 0.44 | 1.54 | 4.45 | 1.30 | 0.42 | 1.16 | 1.92 |
| ECG | 2.23 | 1.35 | 1.56 | 1.62 | 2.23 | 1.32 | 1.55 | 1.63 |
| CG | 0.37 | 0.05 | 1.89 | 4.73 | 0.37 | 0.05 | 2.07 | 6.24 |
| GA | 1.12 | 0.78 | 2.50 | 6.28 | 1.11 | 0.77 | 2.50 | 6.17 |
| CAF | 22.98 | 3.59 | -0.14 | -0.42 | 22.92 | 3.61 | -0.18 | -0.38 |
| Mean | 8.90 | 2.53 | 1.07 | 2.29 | 8.89 | 2.51 | 1.03 | 2.17 |
| SD | 9.19 | 2.60 | 1.04 | 2.43 | 9.18 | 2.57 | 1.05 | 2.43 |

**Table S2.** Mean (SD) of observed and estimated TFCC training and validation data using Y-Net.

| **TFCC** | **Observed data** | | **Estimated data** | |
| --- | --- | --- | --- | --- |
|  | **Training** | **Validation** | **Training** | **Validation** |
| GC | 7.22 | 5.12 | 7.37 | 6.72 |
| ECG | 26.06 | 20.63 | 25.08 | 24.37 |
| C | 3.32 | 2.87 | 3.07 | 2.93 |
| EC | 6.88 | 5.98 | 6.38 | 6.18 |
| EGCG | 20.60 | 18.70 | 20.98 | 19.22 |
| GCG | 1.31 | 1.28 | 1.35 | 1.24 |
| ECG | 2.28 | 2.12 | 2.42 | 2.19 |
| CG | 0.38 | 0.36 | 0.38 | 0.37 |
| GA | 1.06 | 1.23 | 1.14 | 1.25 |
| CAF | 22.33 | 24.34 | 22.54 | 21.54 |
| Mean | 9.14 | 8.26 | 9.07 | 8.60 |
| SD | 9.41 | 8.74 | 9.32 | 8.87 |
